# Predicting eye movements from deep neural network activity decoded from fMRI responses to natural scenes

**DOI:** 10.1101/166421

**Authors:** Thomas P. O’Connell, Marvin M. Chun

## Abstract

Computational models of selective spatial attention can reliably predict eye movements to complex images. However, researchers lack a simple way to measure covert representations of spatial attention in the brain and their link to overt eye movement behavior, especially in response to natural scenes. Here, we predict eye movement patterns from spatial priority maps reconstructed from brain activity measured with functional magnetic resonance imaging (fMRI). First, we define a computational spatial attention model using a deep convolutional neural network (CNN) pre-trained for scene categorization. Next, we decode CNN unit activity from fMRI activity and reconstruct spatial priority maps by applying our computational spatial attention model to decoded CNN activity. Finally, we predict eye movements in a subsequent behavioral experiment within and between individuals using reconstructed spatial priority maps. These results demonstrate that features represented in CNN unit activity can guide spatial attention and eye movements, providing a crucial link between CNN models, brain activity, and behavior.

---

The human visual system is exposed to more information than could possibly be processed. To mitigate this problem, selective spatial attention allows the visual system to modulate perceptual and cognitive processing towards important regions of space^1,2^. Computational modeling of selective spatial attention has been an area of focused research for the past 20 years, and numerous computational hypotheses have been proposed regarding the features, representations, and operations necessary to guide spatial attention (for reviews, see ^3-5^). Central to most selective spatial attention models is a spatial priority map, a representation that tags the priority of different regions in space for the allocation of covert attention and overt attention such as eye movements^6-8^. To compute spatial priority maps, these models have traditionally integrated information from multiple sources, including bottom-up factors such as stimulus-driven visual salience and top-down factors such as volitional control, behavioral relevance, selection history, and reward. The spatial priority maps produced by many of these computational models accurately predict the allocation of overt spatial attention in humans—usually operationalized as eye movement patterns—towards natural and synthetic stimuli^3,4^.

Recently, selective spatial attention models using deep convolutional neural network (CNN) architectures have yielded state-of-the-art prediction of human eye movement patterns (for a review see ^5^). Inspired by the importance of object information in guiding spatial attention, these models extract attention-relevant features from images using CNNs that have been pre-trained to accurately categorize natural object stimuli^9,10^. Kümmerer et al. (2015)^11^ demonstrated that linear combinations of unit activity from a well-known object recognition CNN^9^ can predict eye movement behavior. Further improvements in predicting eye movement patterns have been obtained by using deeper CNNs for feature extraction, non-linear readout layers, and recurrent readout layers^12-15^. The top ten models on the MIT Saliency Benchmark (http://saliency.mit.edu/results_mit300.html), which ranks selective spatial attention models according to their success in predicting eye movement patterns, all rely on some kind of deep neural network architecture. The most predictive model^14^ explains 87% of the variance in eye movement patterns.

Despite this rich computational modeling literature, the features and computations that guide representation of spatial attention in the brain remain unclear. Eye tracking has long provided a reliable, high-resolution method to measure overt spatial attention towards natural stimuli. However, the internal neural representations of covert spatial attention evoked by natural stimuli are less accessible. This limitation impedes efforts to understand the nature of spatial attention representations in the brain and predict eye movement behavior from brain activity. Initial attempts to access internal representations of spatial attention are promising, but not easily adaptable to natural scene stimuli or prediction of eye movements. As a pioneering example, Sprague and Serences (2013)^16^ reconstructed spatial priority maps from multivariate patterns of blood-oxygen-level-dependent (BOLD) activity evoked by spatially-constrained, high-contrast checkerboard patterns, demonstrating that population activity captured in distributed voxel patterns can support topographic representations of the spatial environment. They further demonstrated that such topographic representations are modulated by overt spatial attention, covert spatial attention, and working memory. However, the controlled nature of their stimuli and spatially-constrained decoding model preclude this approach from being applied to natural scene stimuli. Additionally, their approach did not provide a way to link specific computational theories of spatial priority to overt eye movement behavior.

Thus, novel approaches are needed to access and reconstruct high-resolution spatial representations from BOLD activity evoked by natural stimuli and link these representations to spatial attention behaviors. Such techniques would provide insight regarding the features and computations that support spatial attention in the brain and guide attention-related behaviors, such as eye movements, in naturalistic environments. Furthermore, these efforts provide novel ways to assess the biological validity of computational spatial attention models.

Here, we reconstruct representations of spatial attention priority from BOLD activity to predict eye movement behavior in natural scenes. This was possible using a fMRI decoding model that predicts CNN unit activity from BOLD activity. First, we defined a computational model of spatial attention that leverages the representational structure of unit activity in several layers of a CNN pre-trained for scene categorization to predict human eye movement behavior. Next, we used partial-least squares regression (PLSR) to decode multivariate patterns of CNN unit activity from multivariate patterns of BOLD activity. Finally, we reconstructed spatial priority maps from BOLD-decoded CNN activity using the computationally-defined spatial attention model. To behaviorally-validate reconstructed spatial priority maps, we show that a given participant’s reconstructed priority maps predict their own fixation patterns in a subsequent eye tracking experiment. Furthermore, group-average priority map reconstructions predict fixation patterns in a left-out participant. Finally, we externally validate our model and show that group-average priority maps reconstructions from all participants predict fixation patterns measured in an independent set of participants^17^. To our knowledge, this work represents the first instance that representations of covert spatial attention are reconstructed from BOLD activity patterns to predict eye movements to natural scenes. Additionally, these results show that features captured in CNN unit activity map onto representations of spatial attention priority in the human brain and are sufficient to guide eye movements. Overall, we provide a proof-of-concept for a novel approach to assess the biological validity of CNNs and computational spatial attention models, and, more generally, to link complex behavior, computational theory, and brain activity measurements.

## RESULTS

Eleven participants viewed brief presentations (250 ms) of natural scene images while undergoing functional MRI and completing an old/new detection task. Participants were instructed to fixate on a central fixation dot throughout the experiment, and the short presentation time of 250 ms was chosen to ensure that participants did not have time to initiate a saccade while the image was being presented. Significant sensitivity (*d’*) in the detection task was observed across participants (*M* = 2.04, *SEM* = 0.216, *t_10_* = 9.43, *P* = 2.71 x 10^-6^), indicating participants were attentive to the stimuli throughout the experiment. The following day, participants viewed the same images while their eye movements were monitored with an eye tracking camera. Eye movements were recorded in a separate session to ensure that covert spatial representations evoked in the fMRI experiment were not contaminated by co-occurring eye movements. Overall, we aimed to assess a CNN-based computational model of selective spatial attention by using CNN unit activity decoded from BOLD activity evoked during the fMRI session to predict subsequent eye movement behavior outside the scanner. All analyses were run in the following functionally-localized regions of interest (ROIs): V1, V2, V3, hV4, LOC, PPA, FFA, OPA, RSC, IPS, and sPCS.

### Spatial attention model definition

As the foundation for our spatial attention model, we used a goal-directed CNN with a deep architecture^10^ trained for scene categorization^18,19^. This architecture consists of 18 spatially-selective layers that compute alternating convolution and non-linear max-pooling operations (**Fig. 1a**). Representations within these layers are organized along the two spatial dimensions of the image and a feature-based dimension capturing filters (channels) that through learning have become tuned to different visual features that support scene categorization. These representations can be thought of as a series of two dimensional feature maps, each of which shows where a different visual feature is present in an image (**Fig. 1b**). The spatially-selective layers feed into two spatially-invariant fully-connected layers, which in turn provide input to a softmax layer that computes a probability distribution over a set of 365 scene categories.

**Figure 1.**
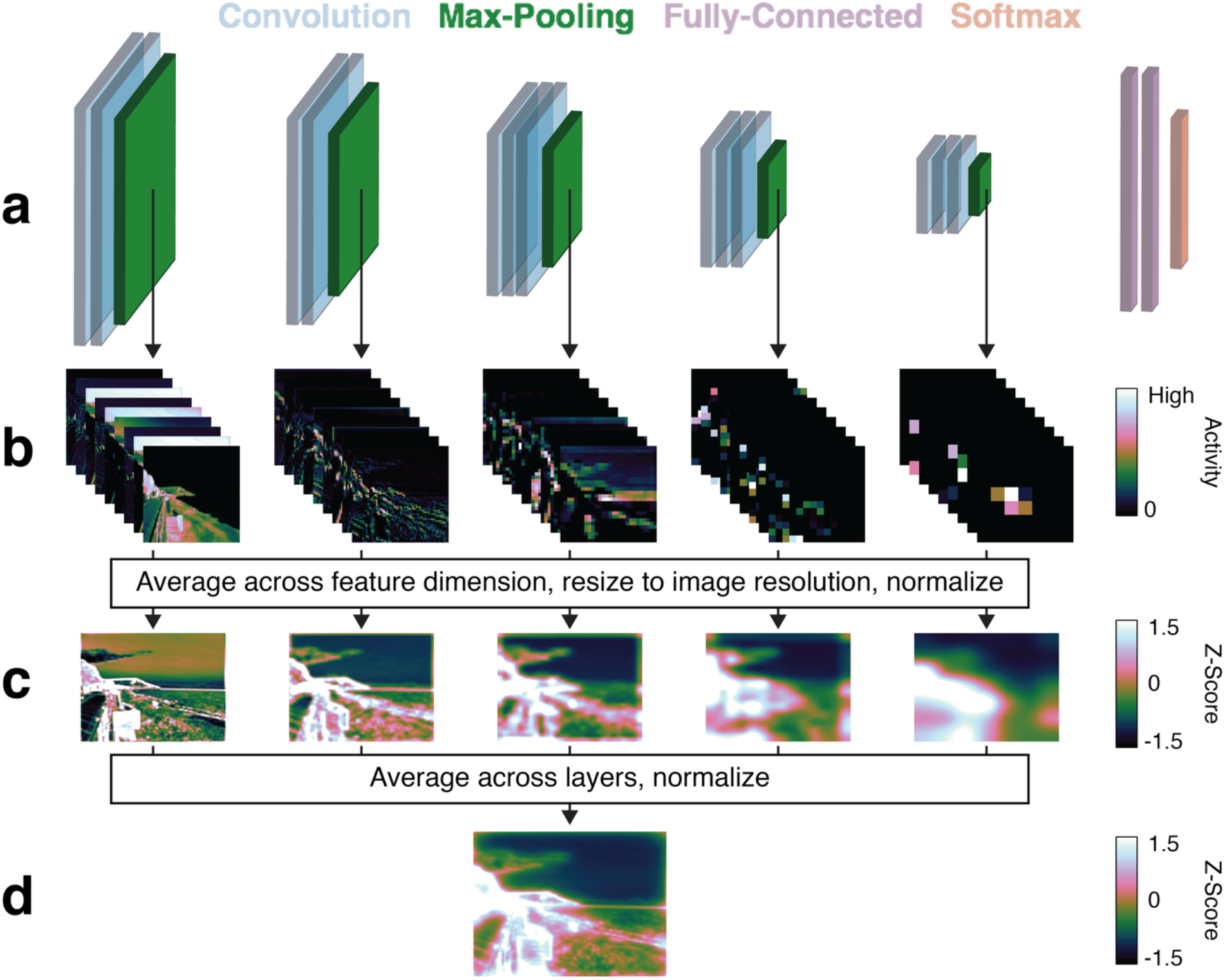
Schematic of the selective spatial attention model. (**a**) A hierarchical CNN trained for scene categorization parametrized each image from pixel-space into a computational feature-space^10,18,19^. (**b**) Unit activity was extracted from the five pooling layers to sample activity from across the CNN hierarchy. Activity in each layer is organized along the x and y spatial dimensions of the image and a feature-based dimension capturing the filter (channel) being applied across the image. This provided a stack of feature maps within each layer demonstrating the location of many different visual features in the image. (**c**) We averaged across the spatial dimension, resized the resultant activity map to the full image resolution (600 x 800 pixels), and normalized each map across spatial locations to have zero mean and unit standard deviation. This process produced a single activity map for each layer showing where in the image visual features captured by filters in the layer were present. (**d**) Finally, we averaged activity maps across layers and renormalized the resultant maps to get computational spatial priority maps for each image. This computational model significantly predicted where individuals from two indenependt data sets allocated overt spatial attention in the images, as measured by eye movements.

Our model of selective spatial attention takes unit activity from the five pooling layers, averages across the feature-based dimension, resizes the resultant maps to the image resolution (800 x 600 pixels), and normalizes values across pixels to zero mean and unit standard deviation to produce a single spatial activity map for each layer that shows where visual features captured by the CNN were present in that layer (**Fig. 1c**). We then average these layer-specific activity maps to produce an overall spatial priority map for each image (**Fig. 1d**). In the common nomenclature of spatial attention modeling, this model represents a bottom-up approach that relies purely on the location of visual features in the image with no top-down weighting of features driven by a volitional control mechanism, selection history, or reward ^20^. This approach of linearly combining unit activity sampled from across the hierarchy in a CNN was inspired by the DeepGaze I spatial attention model^11^.

To assess the goodness of fit between computational spatial priority maps produced by our model and eye movements, we used the Normalized Scanpath Salience (*NSS*) metric, defined as the average value in normalized spatial priority maps at fixated locations^21^. Prior to calculation of *NSS*, spatial priority maps are normalized to have zero mean and unit standard deviation. Thus, *NSS* indicates, in standard deviations, how well a spatial priority map predicts fixated locations within an image. Positive values indicate that fixated regions were accurately predicted and values close to zero indicate that fixated regions were not accurately predicted. The normalization procedure ensures that null prediction will produce expected values of zero, allowing us to parametrically assess significance by comparing *NSS* across participants to zero using a one-sample, two-tailed t-test. We find that spatial priority maps computed by our model produce *NSS* values significantly greater than zero in our primary data set of 11 participants (*M* = 0.803, *SEM* = 0.0841, *t_10_* = 9.54, *P* = 2.45 x 10^-6^) and our external validation data set of 22 participants (*M* = 1.199, *SEM* = 0.0116, *t_21_* = 103.58, *P* = 6.16 x 10^-30^), indicating our spatial attention model significantly predicted eye movement behavior in two independent data sets. Note that the *NSS* values for this model are lower other CNN-based computational spatial attention models on the MIT Saliency Benchmark http://saliency.mit.edu/results_mit300.html). Computational spatial attention models commonly include spatial smoothing and center-bias correction, both of which greatly increase *NSS* values. To facilitate straightforward interpretation of which factors drive significant prediction of eye movements from spatial priority maps reconstructed from fMRI activity, these steps were excluded from our model.

### Reconstruction of spatial priority maps from BOLD activity

We used partial least squares regression (PLSR) to decode patterns of CNN unit activity from patterns of BOLD activity and then reconstructed spatial priority maps from decoded CNN activity (**Fig. 2**). We first reduced the dimensionality of both CNN activity and BOLD activity using separate principal component analyses (PCA). Next, we used PLSR to learn a transformation between BOLD components and CNN components. Precisely, BOLD components were used as predictor variables to estimate CNN components in a leave-one-run-out (LORO) cross-validated manner, such that BOLD components from eleven runs of the fMRI experiment were used as training data to learn the transformation from BOLD components to CNN components. This learned transformation was then applied to BOLD components from the left out run to decode CNN components for each trial in that run. Finally, the decoded CNN components were multiplied by the transpose of the PCA transformation matrix to reconstruct the full space of CNN unit activity. This pipeline was applied separately for each participant, ROI, and CNN pooling layer, producing five sets of BOLD-decoded CNN features for each image in each ROI in each participant (**Fig. 2a**). This decoding approach can be viewed as learning a transformation to express BOLD activity in a common CNN-defined feature space that varies along the two spatial dimensions of the input stimuli and one feature-based dimension.

**Figure 2.**
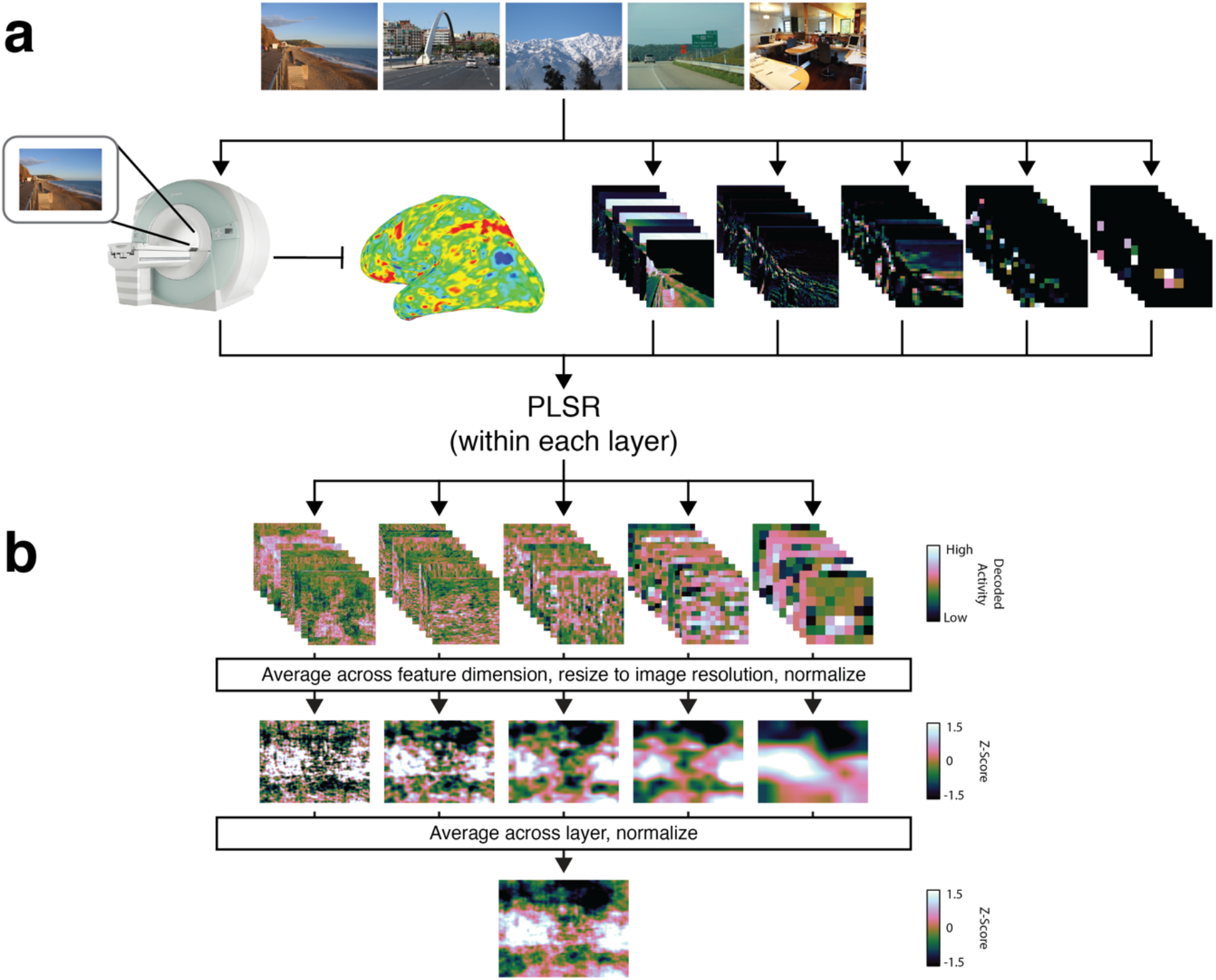
(**a**) fMRI was used to measure evoked BOLD activity for each image in the experiment. Additionally, each image was parameterized into a CNN feature-space. We learned the transformation between BOLD activity and CNN activity separately for each CNN layer using PLSR. These learned transformations were applied to left-out data to decode CNN activity for each layer from BOLD activity. (**b**) To reconstruct spatial priority maps from BOLD-decoded CNN activity, we applied the same spatial attention model to computational CNN activity. Decoded unit activity from each layer was averaged across the feature-based dimension to produce reconstructed layer-specific activity maps, which were then averaged together to produce a reconstructed spatial priority map for each image.

Next, we calculated spatial priority maps from the BOLD-decoded CNN activity using the same spatial attention model defined above (**Fig. 2b**). Within each layer, we averaged BOLD-decoded CNN activity across the feature-based dimension to calculate a layer-specific activity map for each layer. These layer-specific activity maps were then averaged together to reconstruct a final reconstructed spatial priority map for each image. Example reconstructions can be seen in **Fig. 3**.

**Figure 3.**
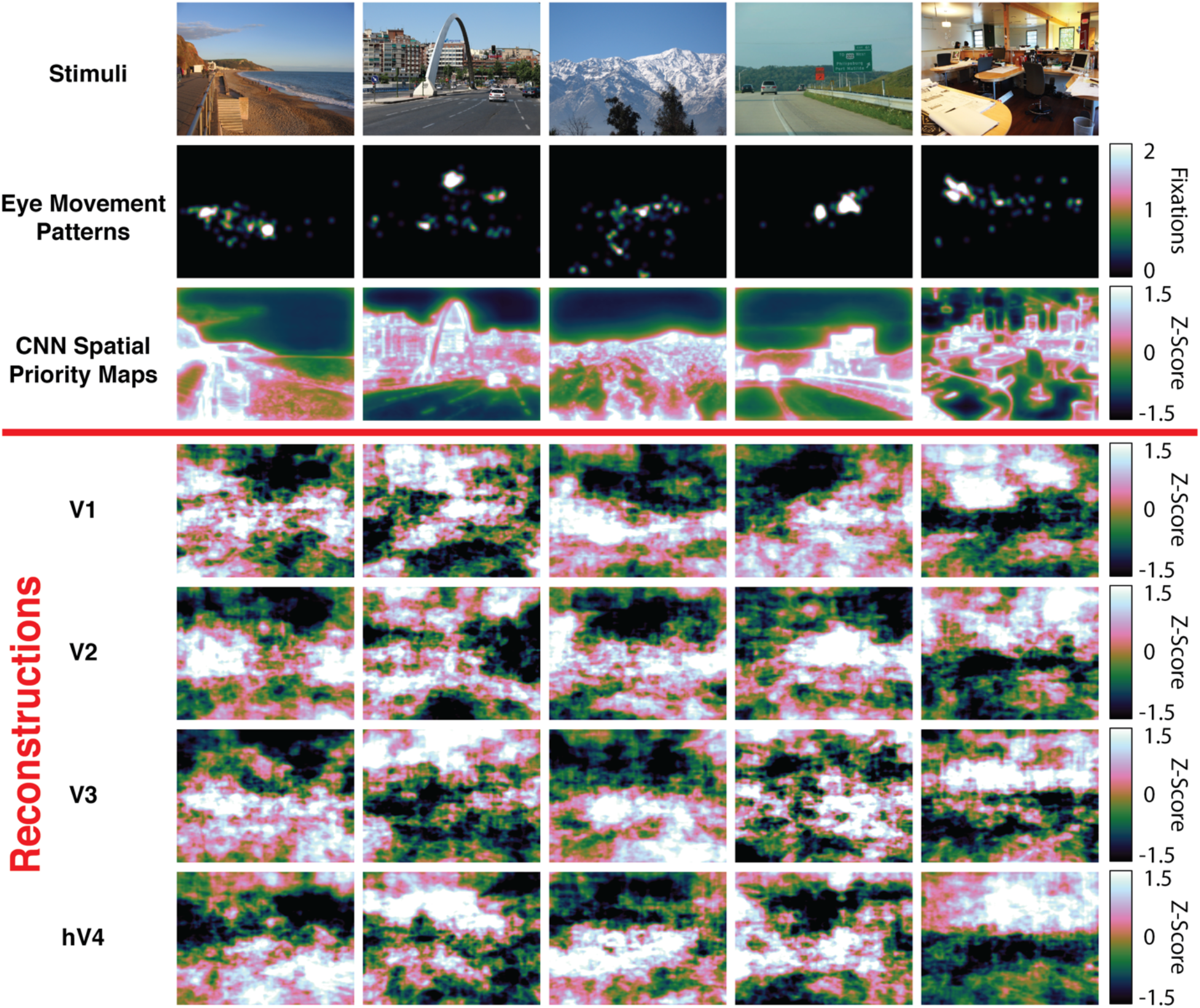
Example stimuli, eye movement patterns, computational spatial priority maps, and reconstructions from early visual ROIs. Each column corresponds to an image that was shown to participants during the fMRI experiment. Fixation maps were averaged across all participants in the external validation data set (*n* = 22) and reconstructions were averaged across all participants in the main experiment (*n* = 11).

### Reconstructed spatial representations predict eye movements

First, we determined whether spatial priority maps reconstructed from BOLD-decoded CNN activity in a given individual predict that same individual’s out-of-scanner eye movement behavior. To assess the goodness of fit between reconstructed spatial priority maps and eye movements, we used *NSS*. *NSS* metrics calculated in this manner for individual participants are independent from one another, allowing us to parametrically assess significance using one-sample, two-tailed t-tests comparing NSS metrics across participants to zero.

We found *NSS* values significantly greater than zero when comparing individuals’ reconstructed spatial priority maps to their own eye movement patterns on the same images the following day, demonstrating that spatial priority maps reconstructed from BOLD-decoded CNN activity predict allocation of overt spatial attention as measured by eye movement patterns within individuals. At a Bonferroni-corrected threshold applied across ROIs (*P* = 4.55 x 10^-3^), *NSS* values were significantly greater than zero for reconstructions from V1 (*M* = 0.0759, *SEM* = 0.0169, *t_10_* = 4.48, *P* = 1.18 x 10^-3^), V2 (*M* = 0.0821, *SEM* = 0.0166, *t_10_* = 4.94, *P* = 5.89 x 10^-4^), V3 (*M* = 0.103, *SEM* = 0.0155, *t_10_* = 6.61, *P* = 6.00 x 10^-5^), and hV4 (*M* = 0.0821, *SEM* = 0.0158, *t_10_* = 5.46, *P* = 2.75 x 10^4^).

**Figure 4.**
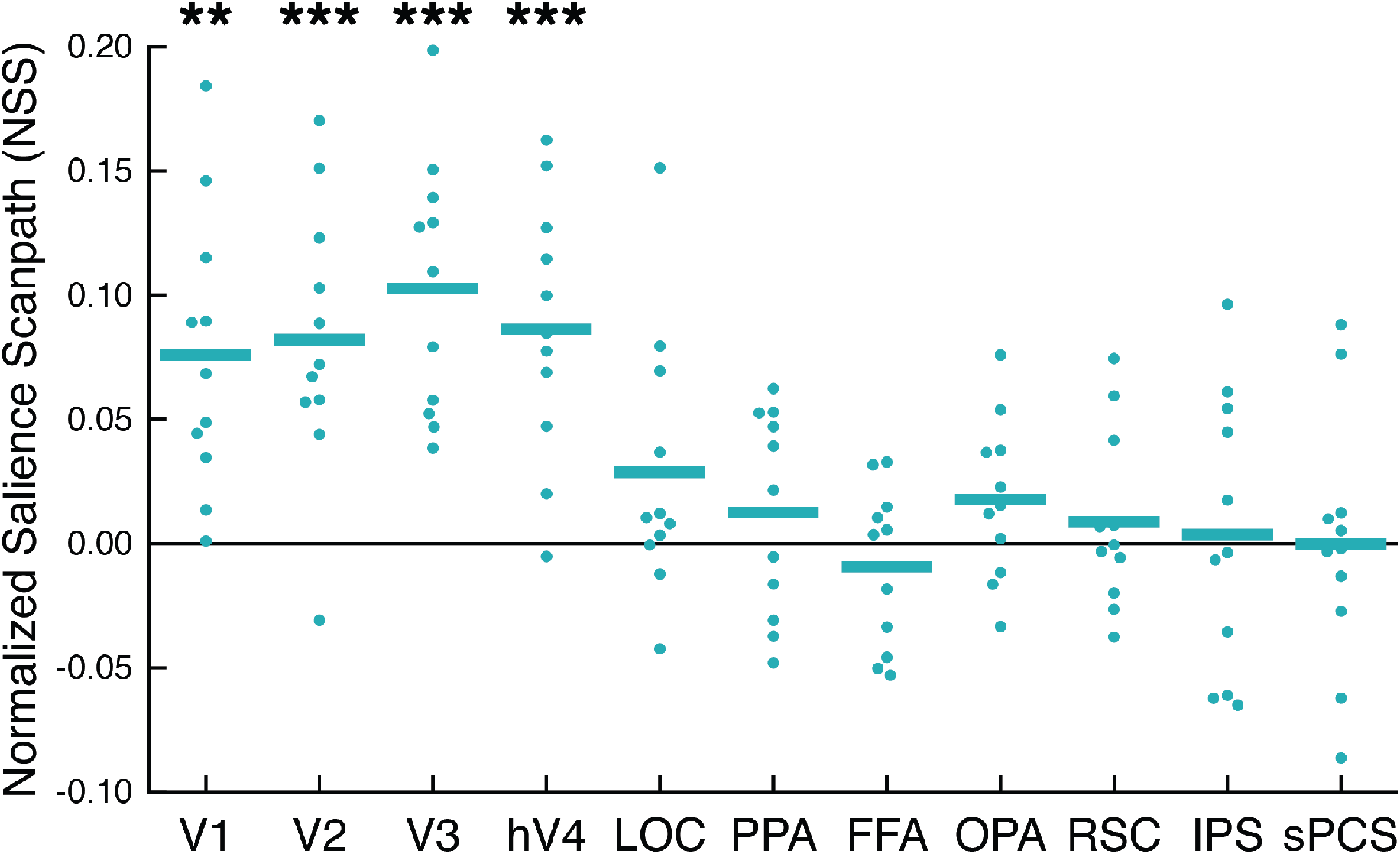
Reconstructed spatial priority maps predicted eye movement patterns within individuals. Eye movements for a given individual were predicated using that individual’s own reconstructions, and significance was determined across participants (*n* = 11) using a one-sample, two-tailed t-test. Positive *NSS* values indicate that reconstructed spatial priority maps predict eye movement patterns. Functional ROIs are plotted along the x-axis, and *NSS* is plotted along the y-axis. Dots represent *NSS* scores from individual participants and bars represent the average *NSS* score across participants. * *P* < 1 x 10^-2^, ** *P* < 4.5 x 10^-3^ (Bonferroni-corrected threshold), *** *P* < 1 x 10^-3^.

### Internal validation: prediction from cross-validated group-average reconstructions

Next, we determined that group-average spatial priority map reconstructions generalize to predict a left-out participant’s eye movements. We decoded the same set of CNN activity from all participants’ BOLD activity; thus, our decoding and reconstruction pipeline can be viewed as an alignment operation that transforms individuals’ BOLD activity from anatomical brain space into a common CNN-defined feature space. This allows for straightforward model-based pooling of brain activity across participants. In a leave-one-subject-out (LOSO) cross validated manner, we computed group-average spatial priority map reconstructions for each image. We then took the left-out participant’s eye movement data and calculated *NSS* to assess how well the group-average reconstruction predicted an unseen participant’s fixation patterns.

Analyses in the leave-one-out folds are not independent; the same participants’ reconstructions are included in the group-averages for multiple folds. This means that parametric assessment of significance using one-sample, two-tailed t-tests may overestimate the degrees of freedom. Therefore, we assessed significance using permutation testing. We randomly shuffled reconstructions with respect to image labels 1,000 times and ran the prediction analysis using permuted sets of reconstructions to generate null distributions of *NSS* values for each ROI. Based on these empirically estimated null distributions, prediction of eye movement patterns using reconstructions from V1 (*M* = 0.179, *SEM* = 0.0101, *P* < 1 x 10^-3^), V2 (*M* = 0.242, *SEM* = 9.62 x 10^-3^, *P* < 1 x 10^-3^), V3 (*M* = 0.271, *SEM* = 0.0102, *P* < 1 x 10^-3^), and hV4 (*M* = 0.218, *SEM* = 0.0100, *P* < 1 x 10^-3^) were significant at the Bonferroni-corrected threshold, consistent with the results observed for prediction within individual participants.

### External validation: prediction from overall group-average reconstructions

As a more rigorous test of generalizability, we used reconstructed spatial priority maps averaged across all participants to predict eye movements made by an independent set of 22 participants who viewed the same images as part of a previous experiment^17^. Overall group-average reconstructed spatial priority maps were calculated using all participants in the fMRI experiment. We calculated *NSS* for each of the 22 participants in the external validation data set to determine how well the group-average reconstruction predicted eye movement behavior in each novel individual.

To maintain consistency with the internal validation analysis, we again assessed prediction significance by estimating empirical null distributions of *NSS* values using permutation testing. Group-average reconstructions were randomly shuffled with respect to image labels 1,000 times, and the prediction analysis was run using these permuted sets of reconstructions to estimate a null distribution for each ROI. Consistent with the internal validation analysis, eye movements in this independent set of participants were significantly predicted by group-average reconstructions from V2 (*M* = 0.269, *SEM* = 7.83, *P* < 1 x 10^-3^), V3 (*M* = 0.279, *SEM* = 8.57 x 10^-3^, *P* < 1 x 10^-3^), and hV4 (*M* = 0.253, *SEM* = 7.57 x 10^-3^, *P* < 1 x 10^-3^). Prediction from reconstructions in V1 (*M* = 0.211, *SEM* = 0.0105, *P* = 5 x 10^-3^) fell just below significance at the Bonferroni-corrected threshold (*P* = 4.55 x 10^-3^), but remained significant at an uncorrected threshold of *P* < 1 x 10^-2^.

**Figure 5.**
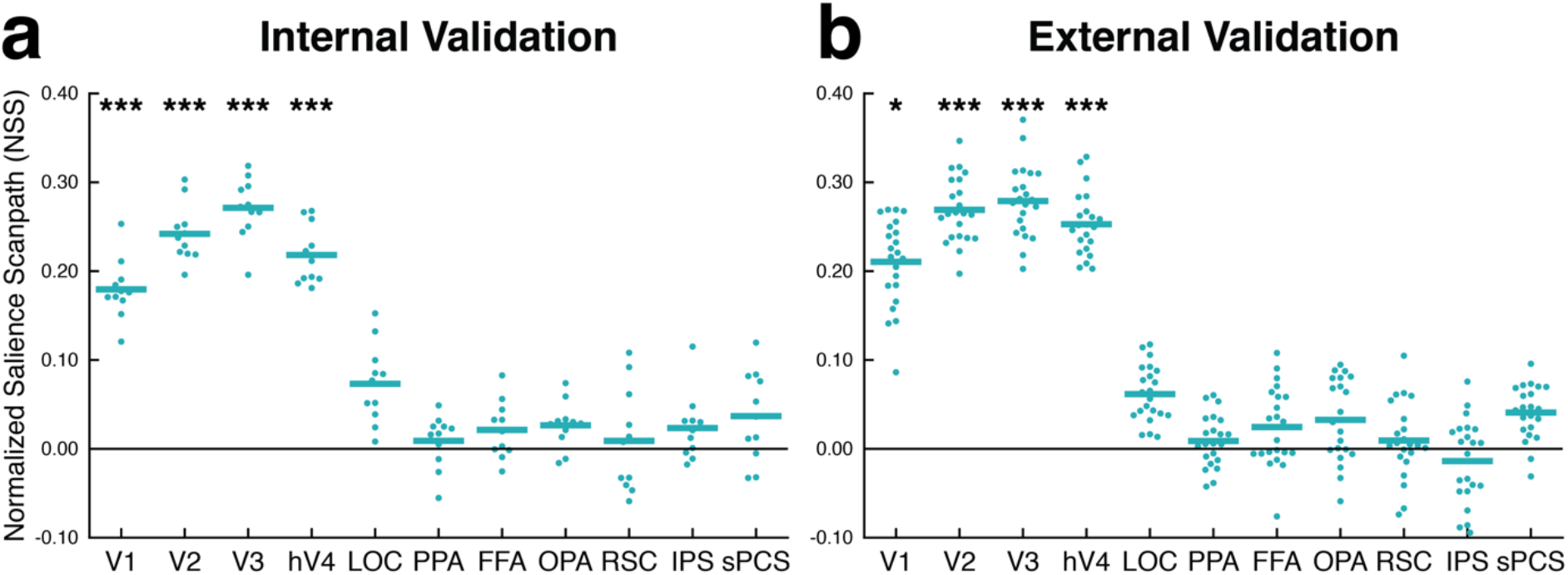
(**a**) Group-average reconstructions predicted eye movements in a left-out participant. Reconstructions were averaged across participants in a leave-one-subject-out cross-validated fashion and used to predict eye movements in the left-out participant. Significance was determined using permutation testing. Dots represent *NSS* scores from individual participants in the main experiment (*n = 11*) and bars represent the average *NSS* score across participants. (**b**) Group-average reconstructions from all participants in the main experiment predicted eye movements from participants in an independent external validation data set^17^. Significance was determined using permutation testing. Dots represent *NSS* scores from individual participants in the external validation data set (*n* = 22) and bars represent the average *NSS* score across participants. * *P* < 1 x 10^-2^, ** *P* < 4.5 x 10^-3^ (Bonferroni-corrected threshold), *** *P* < 1 x 10^-3^.

## DISCUSSION

We reconstructed spatial priority maps from BOLD activity to predict eye movement behavior in natural scenes. This was possible with a novel analysis pipeline to 1). decode CNN unit activity from BOLD activity, 2). reconstruct spatial priority maps from decoded CNN activity using a computational spatial attention model, and 3). predict eye movement patterns from the reconstructed spatial priority maps. Within participants, reconstructed spatial priority maps predicted fixation patterns, demonstrating that features captured in CNN unit activity map onto representations of spatial attention priority in the human brain and are sufficient to guide overt eye movement behavior. Additionally, group-average reconstructed spatial priority maps predicted eye movements in novel individuals both within the current experiment and in an independent validation set, suggesting consistency across individuals in how spatial attention is allocated to natural scenes. These results demonstrate a new approach to access representations of covert spatial attention in the brain and to link these representations to overt eye movement behavior.

### Accessing and reconstructing representations of spatial attention

While eye tracking has long provided a robust and reliable method to measure the allocation of overt spatial attention, methods to access high-resolution representations of covert spatial attention in the brain have not been as readily available. Early fMRI work exploring the effect of spatial attention on BOLD activity in early visual and parietal cortices found univariate signal changes that co-vary with deployment spatial attention to different visual hemifields^22-24^, and such hemifield manipulations have been exploited to find evidence of spatial representations driven by low-level salience^25,26^. However, these hemifield manipulations provide only a very coarse measure of how spatial attention is distributed across the visual field. Multivariate BOLD reconstruction has shown greater promise in accessing high-resolution spatial representations in the brain, successfully reconstructing simple contrast patterns^27,28^, handwritten digits^29^, natural images and movies^30,31^, and faces^32^. More recently, reconstruction techniques have been used to access covert spatial representations in the brain. Sprague and Serences (2013)^16^ showed that spatial representations evoked by low-level checkerboard stimuli could be reconstructed from BOLD activity patterns during allocation of overt attention, covert attention, and working memory maintenance. Sprague et al. (2014)^33^ reconstructed spatial representations from BOLD population activity during working memory maintenance and found an increase in amplitude in the reconstructions at remembered target locations but not forgotten locations. To reconstruct spatial representations, both experiments use an inverse encoding model that quantifies the location of a circular checkerboard stimulus using a series of tiled cosine functions that cover the full stimulus display. While this approach can successfully reconstruct covert spatial representations from controlled stimulus arrays, it is not easily adaptable to reconstruct covert spatial representations evoked by natural stimuli.

Accessing representations of spatial attention to natural scenes and linking these representations to eye movement behavior is an important step to validate computational selective spatial attention models. Eye movements to natural scenes have long provided a crucial measure of behavior to validate computational models of spatial attention to natural scenes^3,4^. Thus, testing the biological validity of computational theories for spatial attention in the human brain requires a model-based procedure for reconstructing spatial priority maps from BOLD activity evoked by natural scenes and predicting eye movements in the same natural scenes from reconstructed spatial priority maps. We provide such an approach, and our work represents a proof-of-concept for an additional measure beyond behavioral prediction of eye movements to adjudicate between different computational spatial attention models.

### Testing the biological validity of CNN features as a model of scene representation for guiding attention and eye movements

The current experiment builds on recent work finding a strong correspondence between representations in goal-directed CNNs trained for visual categorization and brain activity^34-36^. Unit activity in object categorization CNNs predicts neural firing to natural objects in macaque IT and V4^37^. CNNs with better categorization performance also provided superior fits to neural firing patterns. CNN unit activity contains the same representational geometries as object-evoked activity in human and macaque IT^38^. A gradient of similarity to IT geometries was found across CNN layers; deeper layers showed progressively greater similarity to IT geometry. In humans, CNN activity predicts voxel-wise BOLD activity across the ventral stream, and depth of CNN layers corresponded to the gradient of complexity across the ventral stream^39^. Cichy and colleagues (2016)^40^ found evidence for a temporal and spatial hierarchical correspondences between CNNs and brain activity measured via fMRI and MEG. Lower layers of an object categorization CNN contained representational geometries similar to BOLD activity patterns in occipital cortex and MEG activity early in a trial, while higher CNN layers contained geometries similar to those found in temporal and parietal BOLD activity patterns and MEG activity later in a trial. Horikawa and Kamitani (2017)^41^ decoded CNN unit activity from BOLD activity patterns evoked during perception and imagery. They found that decoded unit activity in both tasks captured information about object category, even for novel categories not included in the training set. Additionally, they found that CNN activity decoded from BOLD activity evoked during dreams also contains information about object category^42^.

The current findings move beyond the current literature to show that features represented in CNN unit activity are sufficient to guide spatial attention representation and behavior, which we demonstrate by our computational method that decodes these features from BOLD activity and then combines them according to a computational spatial attention model to predict eye movements. By showing a clear correspondence between BOLD-decoded CNN activity and eye movement behavior, we demonstrate a crucial link between CNNs, brain activity, and behavior that has been absent from the literature. Moving forward, our approach can be applied to assess how features represented in CNN activity might support other behaviors and processes in the brain such as visual categorization or feature-based attention.

### Aligning complex behavior, computational theory, and brain activity measurements

Our technique is a novel approach to assess the biological validity of both CNNs and computational models of spatial attention in the human brain and, more generally, to link complex behavior, computational theory, and brain activity measurements. Over the past decade, two primary approaches have previously been used to assess how well different computational models of neural information processing explain brain activity measurements, including encoding models and representational similarity analysis (RSA). Encoding models predict neural or hemodynamic response patterns from computational model parameters^43,44^. This approach has provided insights in domains including visual image identification^30,31,43^, attentional processing^45,46^, and memory^47^. However, encoding models do not support an easy link to behavior^44^. Representational similarity analysis (RSA) supports comparison of computational model parameters, behavior, and brain activity measurements by computing representational dissimilarity matrices (RDMs) across an entire stimulus set^48-50^. RSA has led to many insights regarding the representations present in the brain to support processing of visual objects, faces, scenes, and other cognitive processes (for a review see ^50^). However, since RDMs must be calculated across an entire stimulus set, RSA does not provide a way to link computational model parameters and brain activity to behavior at the level of individual stimuli.

Given the absence of model-based techniques that allow for stimulus-level prediction of behavior from brain activity measurements, we used a BOLD reconstruction technique to align brain activity into the same representational space as computational spatial attention models and eye movement behavior. In the three-way space of behavior, computational models, and brain activity, behavior and computational models are well aligned. The validation technique for many computational models of neural information processing is prediction of behavior, and subsequently the outputs of computational models often reside in the same space as the behavior of interest. In the domain of overt spatial attention, the behavior of interest (eye movements) reside and are measured in the native dimensions of the stimulus (2D space). The spatial priority maps output by selective spatial attention models reside in the same 2D space as the stimulus, allowing for straightforward comparison between predictions of attentional allocation made by the models and actual eye movement behavior. Given this close alignment between behavior and computational models, one way to solve the three-way alignment problem is to learn a direct mapping between brain activity measurements and computational model parameters. This is the approach we take. The PLSR decoding step in our pipeline can be viewed as an alignment procedure. Instead of framing PLSR as decoding CNN activity from BOLD activity, PLSR can be framed as learning a set of weights that transforms BOLD activity into the same feature-space as CNN activity. In this way, decoding simply reorganizes BOLD activity to vary along the same dimensions along which CNN unit activity varies. Once BOLD activity is expressed in this manner, the same computational attention model applied to computational CNN activity to calculate a spatial priority map can be applied to BOLD activity aligned to the same space as CNN activity. This provides a brain-based spatial priority map that resides in the 2D space as the input stimulus, just like the behavior (here, eye movements) and the computational theory for the representation supporting the behavior (here, a computational spatial priority map). The brain-based spatial priority map can then be used to predict behavior in the same manner the computational model itself is used to predict behavior, allowing for a direct test of how well the brain may support a complex behavior through a theorized model of information processing.

In sum, we demonstrate that high-resolution representations of covert spatial attention can be reconstructed from BOLD activity patterns to predict overt eye movement behavior for natural scene input. These results demonstrate that features represented in CNN unit activity are sufficient to support spatial attention representation and guide eye movement behavior. Beyond this finding, our approach provides a novel technique to assess the link between computational theories of neural information processing, brain activity measurements, and behavior in a unified framework.

## METHODS

### Spatial attention model

*HCNN architecture*. We used Places365-VGG, a variant of the VGG CNN model^10^, trained for scene categorization on the Places365 image set^18,19^. The VGG architecture used in Places365-VGG consists of 21 layers: 13 convolution layers, five pooling layers, and three fully-connected layers^10^. The network takes a 224 x 224 RGB image, with the mean RGB value from the training set subtracted out, as input. The first 18 layers are a series of convolutional layers, consisting of filters with a small receptive field of 3 x 3 and a fixed convolution stride of 1 pixel, followed by spatial max-pooling layers. Units in these layers are organized along three dimensions: the x and y dimensions of the input image and a feature-based dimension that captures which filter (channel) produced the activity for a given feature map. This stack of convolution and pooling layers feeds into the three fully-connected layers. The last fully-connected layer is a softmax classifier that produces a probability distribution over 365 scene category labels.

The Places365 database is a scene-centric large-scale image database consisting of natural scene images downloaded off the internet and labeled across a series experiments by human observers^19^. Places365-VGG was trained on a set of 1,803,460 images and validated on a set of 18,250 images. On a test set of 328,500 images, Places365-VGG produced better classification performance than two other HCNN architectures, AlexNet^9^ and GoogLeNet^51^, trained in the same manner as Places365-VGG on the Places365 image set^19^. Places365-VGG also produced superior classification performance than Places365-AlexNet and Places365-GoogLeNet on four additional scene-centric image sets^19^. Additionally, the filters in Places365-VGG, especially in the later layers, develop receptive fields that detect whole objects^52^. Objects in scenes are known to attract spatial attention^53-55^, suggesting that the spatial features extracted by Places365-VGG should be maximally predictive of spatial attention behavior.

*Spatial priority map calculation*. Each image was resized from its native resolution (800 x 600) to the input size for Places365-VGG (224 x 224) and provided as input to Places365-VGG in Caffe^56^. We limit the features in our model to unit activity from the five pooling layers. After extracting unit activity from each pooling layer, we averaged activity across the feature-based dimension to calculate a single activation map for the layer. The resultant activation map was then resized to full image resolution and normalized across all locations to have zero mean and unit standard deviation. Activation maps were then averaged across layers and re-normalized to zero mean and unit standard deviation to calculate the final spatial priority map.

*Evaluation criteria*. We evaluated the success of our model at predicting human fixation patterns using the Normalized Scanpath Salience (*NSS*) metric^21^. *NSS* is defined as the average salience values at all fixated locations within an image.

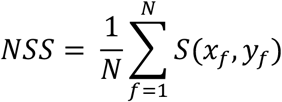

Here, *S* is the spatial priority map, *N* is the total number of fixations, and (*x*_*f*_, *y*_*f*_) is a fixated location.

### fMRI data set

*Participants*. 15 participants from Yale University and the surrounding community underwent fMRI scanning while viewing natural scene images and completing a behavioral old/new recognition task. One participant was excluded because they withdrew from the study early and three participants were excluded for excessive motion, leaving 11 for analysis (6 females, ages 19 -36, mean age = 25.27). Excessive motion for individual runs was defined a-priori as >2mm translation or >3° rotation over the course of the run. A participant was excluded if more than two of their runs showed excessive motion. Nine of the included participants had usable data for all 12 runs, one participant had 11 runs, and one participant had 10 runs. All were right handed and had normal or corrected-to-normal vision. The study was approved by the Yale University Human Subjects Committee. Participant gave written informed consent and were paid for their participation.

*fMRI paradigm and stimuli*. Participants performed an old/new recognition task during scanning. Stimuli were images of natural scenes presented in their native resolution of 800 x 600 pixels. In the scanner, they subtended 17.60 x 13.20 degrees of visual angle. A fixation dot was visible in the center of the display throughout the entire experiment. Participants were instructed to stare at the fixation dot and not move their eyes for the duration of each run. Stimulus presentation and response recording was controlled using Psychtoobox^57^.

Each trial began with a fixation point presented for 1000 ms. Then an image of a natural scene was briefly presented for 250 ms and followed by a 1500 ms response period where participants were asked to indicate via button press (left key = ‘new’, right key = ‘old’) whether they had previously seen the scene in an earlier trial within the run. The short presentation time of 250 ms was chosen to ensure that participants did not have time to initiate a saccade during the image presentation period. After a 1250 ms fixation, participants completed an active-baseline arrow task (5000 ms) where they were asked to indicate what direction a series of four left- or right-facing arrows were pointing. This active-baseline task was followed by a 2000 ms fixation period before the next trial began. Each participant completed 12 runs with 24 trials per run, for a total of 288 trials. Within in each run, 12 scene images were each presented twice, with a lag of 3 to 5 trials between repetitions.

*Imaging and preprocessing*. fMRI data were collected on a 3T Siemens Trio TIM system with a 32-channel head coil at the Yale Magnetic Resonance Research Center. A T1-weighted gradient-echo sequence was used to acquire high-resolution structural images for each participant (TR = 1900 ms, TE = 2.52 ms, flip angle = 9°, FOV = 250 mm, matrix size = 256 x 256, in-plane resolution = 1.0 mm^2^, slice thickness = 1.0 mm, 176 sagittal slices). Functional runs included 3408 task volumes (284 per run) acquired using a multiband gradient-EPI (echo-planar imaging) sequence (TR = 1000 ms, TE = 30 ms, flip angle = 62°, FOV = 210 mm, matrix size = 84 x 84, in-plane resolution = 2.5 mm^2^, slice thickness = 2.5 mm, 51 axial-oblique slices parallel to the ac-pc line, multiband acceleration factor = 3). The first 5 TRs of each functional run were discarded, leaving a total of 3348 task volumes 279 per run).

Data were analyzed using AFNI^58^ and custom scripts in Matlab (R2014a, The MathWorks, Inc., Natick, Massachusetts, United States) and Python (version 2.7, Python Software Foundation). Functional data were despiked, corrected for motion, and aligned to the high-resolution MPRAGE. Cortical surface reconstruction was completed using Freesurfer^59-64^. Functional data was projected from volumetric face to the cortical surface, and all subsequent analyses were done in surface space. In surface space, data were spatially smoothed using a 5 mm full-width, half-maximum Gaussian filter. 12 motion parameters (roll, pitch, yaw, superior displacement, left displacement, posterior displacement, and the derivatives of these parameters) were regressed from the functional data, and the error terms from this regression were used for all subsequent analyses. Functional data for each trial were averaged from TR 5 to TR 8 to extract a single whole-brain activation pattern for each trial.

*Regions of interest*. Functional data to localize ROIs were collected in a separate scanning session from the main experiment. Functional scan parameters for the localizer scans matched the parameters used for the main experiment. Borders of early visual areas (V1, V2, V3, hV4) were delineated on the flattened cortical surface with standard retinotopic mapping techniques^65,66^ using a left/right rotating wedge and expanding/contracting ring of a flickering checkerboard pattern. Five category-specific ROIs were functionally defined using data from two localizer scans ran in a separate MRI session from the main experiment. In each run, participants viewed blocks of rapidly-presented images from the following categories: faces, scenes, objects, and scrambled objects. Scene-selective parahippocampal place area (PPA), retrosplenial cortex (RSC), and occipital place area (OPA) were defined using a [scenes > faces] contrast ^67^1. Object-selective lateral occipital cortex (LOC) was defined using a [objects > scrambled] objects contrast. Face-selective fusiform face area (FFA) was defined using a [faces > scenes] contrast ^68^. Finally, two regions implicated in shifts of attention were functionally localized using data from two additional localizer scans. In these scans, participants alternated between blocks of fixating on a stationary dot in the middle of the screen and blocks of shifting gaze to follow a moving dot that moved to a new random location on the screen every 1000 ms. Attention-selective regions in the intraparietal sulcus (IPS) and superior precentral sulcus (sPCS) were defined using a [shifting fixation > stationary fixation] contrast. Each ROI was defined unilaterally then combined across hemispheres into the bilateral ROIs used for all subsequent analyses.

*Dimensionality reduction*. We reduced the dimensionality of both Places365-VGG unit activity and BOLD activity using principal components analyses (PCA). The number of units in each pooling layer of Places365-VGG are as follows: 802,816 in pool1, 401,408 in pool2, 200,704 in pool3, 100,352 in pool4, and 25,088 in pool5. Within each layer, we start with a [*s* x *u*] matrix, with *s* capturing each scene image (144 total) in the experiment and *u* capturing all units in the layer. Using PCA, we extract a set of 143 component eigenvectors for each layer. This allows us to reduce the original [*s* x *u*] unit-space to a [*s* x *u_PC_*] space, with *u_PC_* capturing 143 principal components (PC) scores across the 143 component eigenvectors. Additionally, the PCAs for each layer produce a [*u* x *u_PC_*] transformation matrix to move between the original unit-space and unit-PC-space. After decoding, the transpose of this transformation matrix will be used to project the *u_PC_* scores decoded from brain activity back into the original Places365-VGG unit-space to reconstruct the full set of unit activity for each layer.

Additionally, separate PCAs were used to reduce the dimensionality of BOLD activity patterns from each ROI. There are different numbers of voxels in each ROI in each participant, and this step normalizes the number of predictor features used for decoding across ROIs and participants. BOLD activity starts in a [*t* x *v*] matrix of voxel activity, with the *t* capturing trials (288 total) and the *v* capturing voxels within a given ROI. We use the same number of components from the Places365-VGG PCAs (143) to reduce BOLD activity patterns to a [*t* x *v_PC_*] space, with *v_PC_* capturing BOLD PC scores across voxels.

*CNN unit activity decoding*. To decode CNN unit activity from the five pooling layers of Places365-VGG from BOLD activity, we used partial least squares regression (PLSR). PLSR is a multivariate machine learning technique that extracts latent variables from a multidimensional input space (here, BOLD activity) and a multidimensional output space (here, Places365-VGG unit activity) then learns a linear transformation between the two sets of latent variables. PLSR has several characteristics that make it ideal for modeling fMRI data^69,70^. First, PLSR works well with data showing high multicollinearity between predictors (here, voxels), as is common for fMRI data. Second, PLSR handles data sets with many more predictors than measurements (here, trials) by reducing the full feature-spaces to a set of latent variables equal in size to the degrees of freedom for the data sets (in this case, number of training trials minus one). Finally, PLSR allows for the leverage of multivariate patterns within both input and output variables, unlike many common fMRI pattern analysis techniques which only leverage multivariate patterns in input variables.

We used PLSR to learn a [*v_PC_* x *u_PC_*] transformation matrix that captures the linear relationship between BOLD components and Places365-VGG unit components. PLSR models were trained in a leave-one-run-out (LORO) cross-validated fashion, such that trials in eleven runs of the fMRI experiment are used as training data to learn the [*v_PC_* x *u_PC_*] weight matrix. We allowed the maximum number of PLSR components (142, equal to the number of BOLD and Places365-VGG components minus one). BOLD components from trials in the left-out run were then multiplied by this learned transformation matrix to decode Places365-VGG components for each trial in the run. These decoded Places365-VGG components were then multiplied by the transpose of the PCA dimensionality reduction matrix to reconstruct the full Places365-VGG unit activity space. The reconstructed Places365-VGG unit activities were then averaged across the two repetitions for a given scene. This approach is analogous to a previously published technique used to reconstruct face images from patterns of BOLD activity (Cowen et al. 2014). Separate PLSR decoders were used to decode Places365-VGG unit activity for each pooling layer layer within each participant from BOLD components extracted from bilateral activity in each of the 11 ROIs.

*Spatial priority map reconstruction*. Spatial priority maps were reconstructed from decoded Places365-VGG activity from all five pooling layers using the same computational spatial attention model defined above. Decoded Places635-VGG unit activity from each pooling layer was averaged across the feature-based filter dimension to produce a single two-dimensional activity map showing for each layer. These activity maps were then resized to the image resolution (600 x 800) and normalized across pixels to have zero mean and unit standard deviation. These layer-specific activity maps were averaged across layers to produce a single reconstructed spatial priority map for each ROI in each participant was then re-normalized to have unit mean and zero standard deviation. These maps were used for all within-participant analyses.

Additionally, to produce group-average reconstructed spatial priority maps, participant-specific reconstructions were averaged across participants. If a given participant was missing BOLD data for a given image, they were excluded from the calculation of the group-average reconstruction for that image. In the internal validation eye movement prediction analysis, these group-average maps were generated using 10 participants (N-1). In the external validation eye movement prediction analysis, these group-average maps were generated using the full set of 11 participants.

*Eye tracking apparatus*. During the follow-up eye tracking session, eye movements were monitored using an Eyelink1000+ eye tracking camera (SR Research, Ottawa, ON, Canada), which uses infrared pupil detection and corneal reflection to track eye movements. The camera was mounted above the participants’ head using a tower setup that stabilized participants’ heads with a chin rest and a forehead rest. Eye movements were recorded monocularly from the participants’ right eyes at 1000 Hz. Participants were positioned 50 cm from an LCD monitor 43 cm diagonal) with a resolution of 1280 x 1024 pixels and refresh rate of 60 Hz. Stimuli were presented in their native resolution of 800 x 600 pixels and subtended 23.99 x 17.98 degrees of visual angle. Stimulus presentation and response recording were controlled using Psychtoolbox^57^.

*Eye tracking paradigm and stimuli*. The day after the fMRI session, participants returned to complete a surprise recognition memory test on the stimuli from the day before while their eye movements were monitored using an eye tracking camera. Half of the stimuli were the 144 scene images from the fMRI experiment and the other half were 144 lure scene images the participants had never seen. They were instructed to freely explore each image then provide a response indicating their confidence that they saw the image during the fMRI experiment the preceding day (1 = Defintely old, 2 = Probably old, 3 = Probably new, 4 = Defintely new).

At the beginning of each block, the eye tracking camera was calibrated and validated using a nine-point fixation sequence. Each trial was preceded by a drift check in which the participant stared at a centrally located fixation dot. The camera was re-calibrated in the middle of the block if the spatial error during the drift check on the preceding trial exceeded 1° of visual angle. The mean spatial error across all calibrations and participants was 0.46° of visual angle (SD = 0.10°). After the drift check, an image was presented for 3000 ms and participants’ eye movements were monitored while they explored the images. After 3000 ms, the response scale appeared on the scale and the participants had as much time as they needed to make their memory judgment. After their response was recorded, a new trial began. Overall participants completed 12 blocks of 24 trials, with 12 target images from the fMRI experiment and 12 novel lure images in each block.

*Eye movement prediction*. All fixations prior to 200 ms were discarded to remove the initial centrally located fixation recorded for each trial. Fixations from 300 ms to 2000 ms were included in all eye movement analyses.

For the within-individual validation eye movement prediction analysis, *NSS* was calculated within each ROI for each image using an individual’s own reconstructed spatial priority maps and eye movements on the same image. *NSS* was averaged across all images to compute a single *NSS* metric for each ROI in each participant. Within each ROI, *NSS* across participants was compared to zero using a one-sample two-tailed t-test.

For the internal validation eye movement prediction analysis, *NSS* was calculated with each ROI for each image using a cross-validated group-average reconstructed spatial priority map and eye movements from a left-participant. *NSS* was averaged across images within each ROI and individual, and again compared to zero with a one-sample two-tailed t-test.

### External validation eye movement data set

*Participants*. 22 participants (9 female, ages 18-41, mean age = 20.2) viewed the same images as part of a previously published study^17^. All participants had normal or corrected to normal vision and received partial course credit in exchange for their participation. Each participant provided written informed consent and the study was approved by The Ohio State University Independent Review Board.

*Apparatus*. Eye movements were monitored using an Eyelink1000 eye tracking camera (SR Research, Ottawa, ON, Canada), which uses the same method of infrared pupil detection and corneal reflection to track eye movements as the Eyelink1000+ camera used in the current experiment. Eye movements were monitored monocularly from each participant’s dominant eye at 1000 Hz. As in the current experiment, the camera was mounted above the participants’ heads using a tower setup, the participants’ heads were stabilized with chin and forehead rests, calibration and validation were completed using a nine-point fixation sequence, drift checks were performed before each trial, and the camera was recalibrated at the beginning of every block or if the error exceeded 1° of visual angle during a drift check. The average spatial error across all calibrations and participants was 0.49° of visual angle (SD = 0.13°).

*Paradigm and stimuli*. For detailed methods, see Experiment 1 in O’Connell & Walther (2015)^17^. Participants viewed each image used in the fMRI portion of the current experiment, plus an additional 54 images not included in the current experiment. Participants freely explored each image for 2000 ms to 8000 ms and answered a multiple choice or true/false question about the image after 33% of the trials to ensure attention throughout the experiment.

*Eye movement prediction*. The same fixation selection criteria applied in the current experiment were applied to eye movement data from the external validation data set. Any fixation made in the first 200 ms was discarded, and fixations intiated between 200 ms and 2000 ms were included in subsequent analysis.

For the external validation eye movement prediction analysis, we aimed to predict eye movements from the 22 participants in this data set using the overall group-average reconstructed spatial priority maps from all 11 participants in the current experiment. We calculated *NSS* for each of the 22 participants on each image using reconstructed priority maps from each ROI and averaged across all image to get average *NSS* for each ROI in each participant. Within each ROI, *NSS* across participants was compared to chance using one-sample t-test.

### Data and code availability

The scene-categorization convolutional neural network is available online (http://places2.csail.mit.edu/download.html). The computational spatial attention model and decoding models were implemented in Matlab. This code is available from the authors upon request. The code to calculate NSS metrics is available from the MIT Saliency Benchmark page (http://saliency.mit.edu/results_mit300.html). Beeswarm plots were generated using an application available online http://perceptionexperiments.net/SDU/VC/BeeswarmTool). The data that support the findings in this manuscript are available from the corresponding author upon reasonable request. The data are not publicly available due to restrictions related to participant privacy and consent.

## ACKNOWLEDGMENTS

This work was supported by the Yale FAS Program funded by the Office of the Provost and the Department of Psychology. T.P.O is supported by a US National Science Foundation Graduate Research Fellowship.

## AUTHOR CONTRIBUTIONS

T.P.O and M.M.C. conceived of and designed the study. T.P.O. developed the computational model, decoding methodology, and prediction methodology, collected and pre-processed the data, and analyzed the data. T.P.O wrote the paper with contributions from M.M.C.

